# Risk assessment for the establishment of the Asian giant hornet (*Vespa mandarinia*) in the Pacific Northwest

**DOI:** 10.1101/2021.02.01.429186

**Authors:** Erik D. Norderud, Scott L. Powell, Robert K. D. Peterson

## Abstract

The recent introduction of the Asian giant hornet (*Vespa mandarinia* Smith) in the United States in late 2019 has raised concerns about its establishment in the Pacific Northwest and its potential deleterious effects on honey bees and their pollination services in the region. Therefore, we conducted a risk assessment of the establishment of *V. mandarinia* in Washington and Oregon on a county-by-county basis. Our tier-1 qualitative and semi-quantitative risk assessment relied on the biological requirements and ecological relationships of *V. mandarinia* in the environments of the Pacific Northwest. We based the risk characterization on climate and habitat suitability estimates for *V. mandarinia* queens to overwinter and colonize nests, density and distribution of apiaries, and locations of major human-mediated introduction pathways that may increase establishment of the hornet in the counties of Washington and Oregon. Our results suggest that 5 counties in the region could be at low risk, 48 at medium risk, and 22 at high risk of establishment. For Washington, counties at high risk included Clallam, Clark, Cowlitz, Grays Harbor, King, Pierce, Skagit, Snohomish, and Whatcom. The high risk Oregon counties included Benton, Clackamas, Clatsop, Coos, Douglas, Hood River, Lane, Lincoln, Linn, Marion, Multnomah, Polk, and Yamhill. Many of the western counties of both Washington and Oregon were estimated to be at the highest risk of establishment mainly due to their suitable climate for queens to overwinter, dense forest biomass for nest colonization, and proximity to major port and freight hubs in the region. Considering its negative effects, these counties should be prioritized in ongoing monitoring and eradication efforts of *V. mandarinia*.

## Introduction

Risk assessments are often used to frame the potential impacts that invasive species pose to an ecosystem, economic sectors and industry, and human health and safety. The recent accidental introduction of the Asian giant hornet (*Vespa mandarinia* Smith) is one such invasive species which poses risks to each of those categories in the U.S. *Vespa mandarinia* is the largest hornet species in the world and is a member of the genus *Vespa*, a primary predator of honey bees (Smith-Pardo et al. 2020). The insect was first detected within the U.S. in late 2019 in Whatcom County in northern Washington State, and again in May 2020 in the same county, indicating the possibility of wider spread establishment than just a chance introduction and detection. Further confirmed sightings since May 2020 have prompted federal and state agricultural officials to initiate eradication programs for the pest due to the insect’s propensity to decimate honey bee populations and impact human safety due to its stings, making the insect a threat to the ecosystems, agricultural sectors, and human health and safety within the surrounding areas of their nesting sites.

Biological invasions can be ecologically and economically damaging phenomena which occur in environments around the world. The introduction and establishment of non-native species may disrupt native flora and fauna and their ecosystems, and concomitantly may cause deleterious consequences to a host of economic sectors and at times even public health and safety. To characterize these consequences, risk assessments are regularly developed to frame the problem and ultimately to confer the degree of risk to regulatory agencies and industry pertaining to the establishment of the particular invading species. The type of risk assessment used is ultimately dependent on the data available for a particular stressor. In cases of a new introduction of an invasive species, where an invasion is in its early onset, tier-1 qualitative and semi-quantitative based risk assessments are often employed as a direct result of a lack of quantitative and distribution data for the biological invader in question (Soliman et al. 2014). Tier-1 risk assessments are characterized by deliberate conservative assumptions that represent overestimates of effect and exposure so that the resulting assessment will be conservative and err on the side of safety (SETAC 1994).

With these risks in mind, it is critical to properly frame them to be able to effectively mitigate these hazards to avoid deleterious results to the environment and economy. Therefore, we performed a risk assessment of the establishment of *V. mandarinia* in the Pacific Northwest, focusing on Washington and Oregon.

## Methods

### Problem formulation

The first step of any risk assessment should begin with the initial problem formulation. The problem formulation sets the stage in terms of the scope, steps, and methods of the risk assessment, delineating the ‘stressor’ and its ‘effects’ at its center of focus. In the case of our risk assessment, that stressor is *V. mandarinia* in the U.S. Pacific Northwest and its deleterious effects on the region’s ecosystems and economy. Accordingly, our risk assessment begins with the known biological and ecological characteristics of the stressor, *V. mandarinia*. Additionally, the stressor description also classifies the effect that *V. mandarinia* has on its surrounding ecosystems, focusing on risks to honey bee populations and apiaries. We analyzed the extent of these effects to assess the degree of exposure to these risks in the effects and exposure assessment section of the risk assessment, which primarily analyzed climate and habitat suitability for the insect, factors influencing introduction, and risk to honey bee populations. The final section of our risk assessment drew from the findings of the previous steps, and we ultimately characterized the risks of the establishment of the *V. mandarinia* in the Pacific Northwest using a risk-rating system.

### Stressor description

The Asian giant hornet is prevalent throughout Asia, with its range extending from mainland Asia into Taiwan, Japan, and South Korea (Archer 1995). The insect is in the Vespidae family, within the order Hymenoptera. *Vespa mandarinia* is the largest known species of hornet in the world, ranging from 38-50 mm in length. Although it has a yellow and black abdomen common to many other wasp species, its large orange head and other largely exaggerated facial features make it distinguishable from its close relatives (Lee 2010, Matsuura and Sakagami 1973).

*Vespa mandarinia* utilize a caste system made up of queens, workers, and males, each fulfilling duties integral to the success of the colony (Archer 1995). The life cycle begins with a queen initiating nest foundation after overwintering in a self-excavated cavity in a soft ground-based substrate (Archer 1995). Nest formation takes place over a number of weeks in the late spring. During this period, the queen develops the nest, collects and feeds on arthropods and sap, and prepares to lay eggs (Archer 1995). The colony begins in summer as the queen takes care of her brood and workers eventually begin to emerge. Once the queen has produced enough workers, the duties of the colony are transferred solely to the workers, while the queen remains in the confines of the nest and continues to lay eggs (Archer 1995, Matsuura and Sakagami 1973, Takahashi et al. 2004).

Mating season for *V. mandarinia* begins in early fall, with both new queens and reproductive males emerging (Archer 1995, Matsuura and Sakagami 1973). Males leave the nest before the queens to forage and will need to wait to mate with the newly emerging queens (Matsuura 1984). The activity of the colony gradually decreases in the late fall before ceasing in the early winter, when queens need to find a site to overwinter (Archer 1995). Maturity from egg to adult is approximately 40 days (Matsuura 1984) and the colony cycle lasts approximately 6 months, with the males and workers living for approximately 3 weeks, while queens live as long as 12 months when accounting for their overwintering period (Archer 1995).

The nests are assembled primarily in pre-existing ground-based cavities such as burrows, snake holes, or rotting tree roots (Archer 1995). The nests can be fairly complex and vary in size, with the average containing a few thousand individual cells made from foraged wood-based fibers. A larva matures in each cell (Matsuura and Yamane 1990). Although the average nest can contain a few thousand separate cells, the actual colony size produced from those cells is usually much smaller. Despite this, Archer (1995) observed that a colony produced an average of approximately 200 males and 200 queens in a given cycle in addition to hundreds of workers.

Like other species, *V. mandarinia* has specific habitat preferences and ecological niche requirements which ultimately inform relationships between other species in its surrounding environment and ecosystem. In its native range in parts of mainland and eastern Asia, its distribution has been primarily linked to certain abiotic and biotic factors. Chiefly, it seems to be sensitive to high temperatures and prefers more temperate climates, areas of low elevation and high amounts of precipitation for its nesting site (Alainz et al. 2020, Kim et al. 2020, Zhu et al. 2020). However, there are reports of *V. mandarinia* attacking honey bee colonies at high altitudes, such as in the Himalayan ranges (Batra 1996). Furthermore, queens prefer ‘green’ environments, such as forested areas, parks, agricultural zones, and other herbaceous settings (Kim et al. 2020, Alainz et al. 2020). This finding raises concerns about the risks to wild and cultivated bee populations that are in these environments. In addition, nest colonization within urban greenspaces has the potential to result in human conflicts with *V. mandarinia*, which is a risk to human health and safety. Liu et al. (2016) reported that in only a three month period (July-October when the species is typically active), 42 people died and approximately 1,700 people were injured from suffering multiple stings in China’s Shaanxi Province.

Once *V. mandarinia* has occupied its new environment after initial nest colonization, it must feed and forage. It is a strong flyer has a number of food sources to seek out, most of which are ideally located within a few kilometers of its nesting site (Matsuura and Yamane 1990).

Although the species is most known for its predation on social bees, Matsurra (1984) reported that *V. mandarinia* feed on sap and fruit from a number of different plant species which may lead to crop damage. In its endemic range, queens initially begin feeding on tree sap sources such as *Quercus* (oak) species in mid-late April (Matsurra 1984, Makino 2016). In addition, *V. mandarinia*, has also been observed to be the competitively dominant species among a number of major diurnal sap-feeding species (Yoshimoto and Nishida 2009).

In addition to feeding on plants, *V. mandarinia* is well known to aggressively prey on insect species. The insect attacks beetles, spiders, other social wasp species, but is most well-known for its mass attacks on honey bee species and their colonies. (Matsuura and Yamane 1990, Matsuura 1984). *Vespa mandarina’s* propensity to hunt honey bees has caused issues worldwide (Matsuura and Sakagami 1973), especially in instances where it has become established and local honey bee populations have not had the chance to adapt to its attacks.

Introduction of nonnative species to new territories through natural dispersal such as flying or foraging is unlikely to occur over large geographic distances. However, unlike natural methods of nonnative species introduction into new environments, human mediated introduction is considered the leading cause of nonnative biological invasions, not only in the U.S., but also around the world (Vitousek et al. 1997). This is a result of extensive land transformations producing favorable conditions for invasion, and accidental introductions due to international export and import commercial trade (Vitousek et al. 1997). In the case of *V. mandarina*, accidental introduction to the U.S. in northern Washington, human-mediated introduction through economic trade is the likely reason considering the species was found close to the U.S./Canadian border near ports of entry, and that the region serves as a destination for commercial trade commodities from Asia (Wilson et al. 2020) (Figure 1). This is supported by the captures of *V. mandarinia* in both Vancouver, British Columbia and Washington that were found to originate from two separate lineages (Wilson et al. 2020). The individual captured in British Columbia had DNA from a lineage in Yamaguchi, Japan, while the captured specimen in Washington had DNA linked to a maternal lineage in Chungcheonuk-do, South Korea. Both these introductions in British Columbia and Washington were likely from separate mated queens, although the data could not determine whether these specimens were from the same populations or were introduced at the same time (Wilson et al. 2020). Despite this, the data seem to substantiate the role that human-mediated transport through economic trade has played in the introduction of *V. mandarina* to North America and the Pacific Northwest.

**Figure 1:**
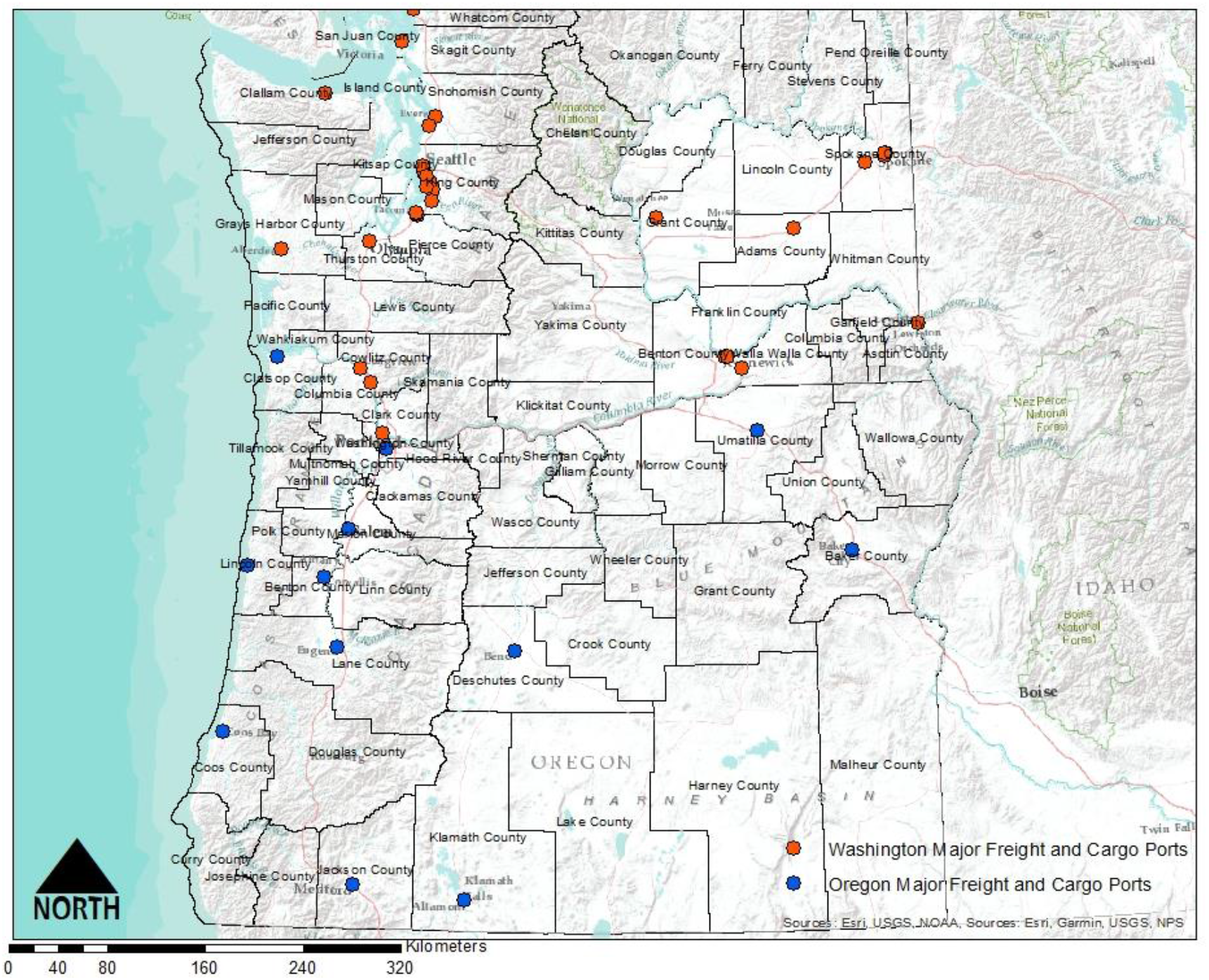
Major freight and cargo ports in the Pacific Northwest. The recent introduction of *V. mandarinia* in North America is thought to have resulted from economic trade activities between the U.S. and Asia.

Beyond global economic trade, cultural pursuits may serve as potential pathways for the introduction of *V. mandarinia*. The species is considered a delicacy in its endemic ranges in Asia, and pupae and adults alike are consumed as food in multiple dishes, and are often semi-domesticated for these purposes (Mozhui et al. 2020). Most ports of entry within the U.S. have safeguarding and inspection measures in place to prevent the importation of live insects and foreign species in cases like this. However, a 100% interception rate is highly unlikely, so the transportation of the insect for cultural purposes should be viewed as a viable potential pathway for its introduction to nonnative regions of the U.S.

### Effects assessment

The effects assessment in our risk assessment relies on *V. mandarinia* queens surviving their overwintering period and successfully initiating a new nest and establishing a colony, potentially causing negative effects to environments in the Pacific Northwest. Those potential deleterious effects occur primarily through feeding and predation strategies of the colony that result in crop damage and attacks on honey bee populations which can thereby impact pollination services.

There are distinct phases that *V. mandarinia* exhibit to attack a honeybee colony. The first phase begins with a solitary scout chemically marking a bee colony or hive by rubbing her terminal gastrite sternite directly on the targeted hive to signal to the rest of the colony of the availability of a source of food (Ono et al. 1995). Once the chemical pheromone has alerted other members of the colony, they will gather en masse and kill the adults in the hive (Matsuura 1984, Ono et al. 1995).

Honey bee species such as *Apis cerana japonica* that have coevolved with *V. mandarinia* in their native ranges have the ability to defend themselves against attack by alerting nestmates of incoming attack using chemical cues (Fujiwara et al. 2018). They use a defense mechanism termed a ‘hot defensive bee ball’ in which hundreds of bees swarm a single hornet and generate enough heat and carbon dioxide around the attacker to kill it (Sugaharo and Sakamoto 2009). Ono et al. (1995) observed through thermal imagery, that the hot defensive bee ball was more than 47 °C (116 °F). Similarly, *A. c. japonica* has been documented to smear both plant based materials and animal feces around hive entrances to disrupt attacks by *V. mandarinia* and the closely related species *Vespa soror* (Matilla et al. 2020).

Once a honey bee colony’s defenses are largely overcome, *V. mandarinia* begins its occupation phase and feeds on the colony’s brood for several days (Ono et al. 1995). For honey bee species that have not coevolved with *V. mandarinia*, such as *A. mellifera*, that only have less effective stingers as a defense (Ugajin et al. 2012), complete annihilation of the colony is a likely outcome when attacked en masse by the hornet, termed the ‘slaughter’ phase (Matsuura and Sakagami 1973, Matsuura 1988). The slaughter phase involves mass attack in which the hornets can quickly dispatch an entire colony, mostly through decapitation using their mandibles. The slaughter event lasts between one and six hours and can result in the deaths of thousands of bees or entire colonies, in which the decapitated bees are often left in massive piles inside the hive (Matsurra 1984, Matsuura and Yamane 1990). The occupation and slaughter phase make the hornet a significant risk to vulnerable non-coevolved bee species. Should *V. mandarinia* become established in ecosystems outside its endemic range, wild bee colonies and apiaries may suffer extremely heavy losses resulting in substantial economic consequences to apiarists and the pollination services provided by wild bee and cultivated honey bees to hundreds of agricultural crops and plant species.

One species of honey bee that may be at potential risk from *V. mandarinia* attack is the European honey bee (*Apis mellifera*). It is a critically important pollinator around the world. The honey bee pollinates hundreds of crop species within the U.S. Honey bees are the foremost insect pollinators, and constitute an estimated economic benefit of nearly $12 billion, or roughly 80% of the total pollination value in the U.S. (Choi and Kwan 2012).

The Pacific Northwest (primarily Washington and Oregon) is the nation’s leader in specialty crops including various varieties of fruits, nuts, and berries, with a total economic value of $4 billion annually (Houston et al. 2018). Considering that the majority of these crops are likely dependent on the pollination services provided by *A. mellifera*, the establishment and naturalization of *V. mandarinia* in the Pacific Northwest could pose high risks for agricultural producers. Beyond agricultural crop varieties, apiculture is also an agricultural sector at risk from the establishment *of V. mandarinia* in the Pacific Northwest. Furthermore, a recent survey of total honey bee colonies within the two states revealed that in June 2020 Washington State had an estimated 114,000 honeybee colonies, while Oregon had an estimated 95,000 (USDA-NASS 2020).

Beyond the potential risks *V. mandarinia* poses to agricultural and apicultural sectors, the insect also poses a risk to health and human safety. The U.S. Census Bureau estimated a population of more than 7.5 million residents in Washington and 4.2 million residents in Oregon in 2019 (U.S. Census Bureau 2019). Although it is statistically unlikely that even a small percentage of those populations would ever interact with *V. mandarinia*, the hornet kills dozens of people per year on average in Japan and causes sting-related injuries to thousands more (New York Times 2020).

### Exposure assessment

The exposure assessment phase of any risk assessment involves drawing upon information from the stressor description and effects assessment and applies relevant data to the environment or ecosystems in question for the purposes of analysis to estimate the degree of risk, impacts, or potential consequences that the stressor may have in those environments. Accordingly, drawing from our stressor description and effects assessment of *V. mandarinia*, our exposure assessment relied on combining its ecology and comparing it to the ecosystems and environments of the Pacific Northwest.

Our analysis primarily focused on regions that match *V. mandarinia’*s climate and habitat suitability requirements and the presence and density of honey bee colonies. Suitable climate for *V. mandarinia* was based on minimum and maximum temperatures using Plant Hardiness Zone Maps of Washington and Oregon and comparing them to the climate in its native ranges in Asia. This is because minimum and maximum temperatures have been cited as necessary abiotic factors critical to the establishment of viable insect populations (Zhu et al. 2020). Therefore, this serves as a good predictor of whether *V. mandarinia* queens would be able to survive their overwintering period.

The U.S. is divided into 13 separate Plant Hardiness zones across 10 °F differences, which are based on minimum winter temperatures. These zones are further classified into two separate zones (A or B) by 5 °F differences. Washington’s Plant Hardiness Zones range from 4A (−34 to -31 °C or -30 to -25 °F) to 9A (−6 to -3.8 °C or 20 to 25 °F). Oregon shares similar zone ratings, which ranges from 4B (−31 to -28.9 °C or -25 to - 20 °F) to 9B (−3.9 to -1.11 °C or 25 to 30 °F) (USDA Agricultural Research Service 2020).

*Vespa mandarinia’*s native ranges in Eastern and Southeast Asia include Plant Hardiness Zones of 6A-13B (Magarey et al. 2008). However, without thorough and up-to-date distribution data of the insect within its endemic ranges or those regions within the Pacific Northwest, it is difficult to pinpoint the precise Plant Hardiness Zones that the insect favors, resulting in what is most likely a highly generalized estimate (Magarey et al. 2008).

Consequently, there is overlap with Plant Hardiness Zones between *V. mandarinia’*s natural range and areas in Washington and Oregon, primarily in the western and coastal regions of each state, but also in some inland regions as well (Figures 2 & 3). Thus, there is appreciable risk that queens may be able to survive their overwintering period within these regions.

**Figure 2:**
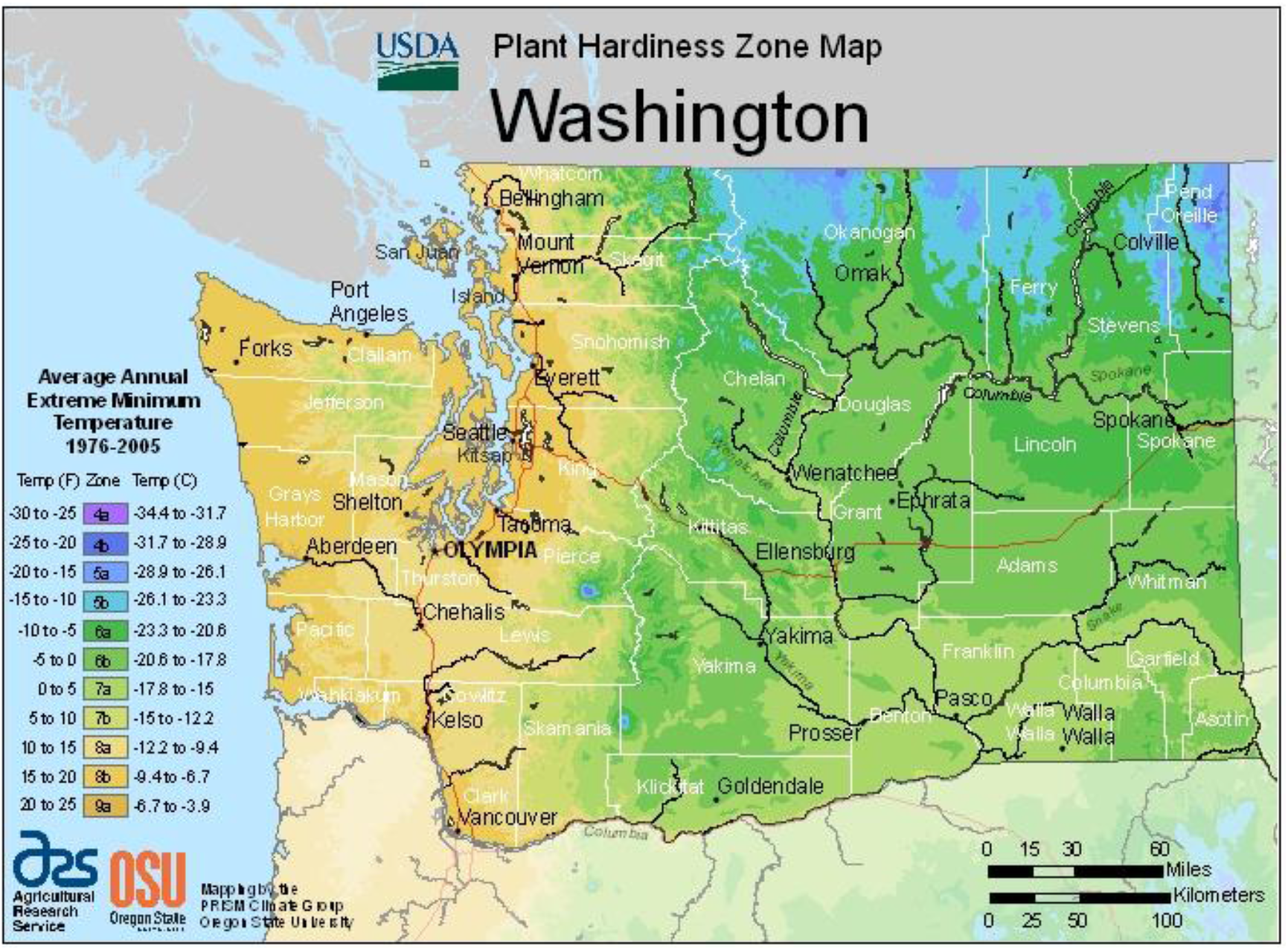
Plant Hardiness Zone (PHZ) Map of Washington State. PHZs are based on the average minimum winter temperature across a 30 year time frame for a region and are used to help growers determine which plants may grow best depending on the zone they inhabit. This risk assessment used PHZs to determine suitable climate where *Vespa mandarinia* may overwinter. Washington PHZs shares some of the same PHZs which are present in *V. mandarinia’*s native ranges. These zones include 6A-9A, which include the majority of Washington State and its apiaries, indicating suitable climate for *V. mandarinia* to overwinter in and predate on honey bee populations. (Source: USDA Agricultural Research Service, 2020)

**Figure 3:**
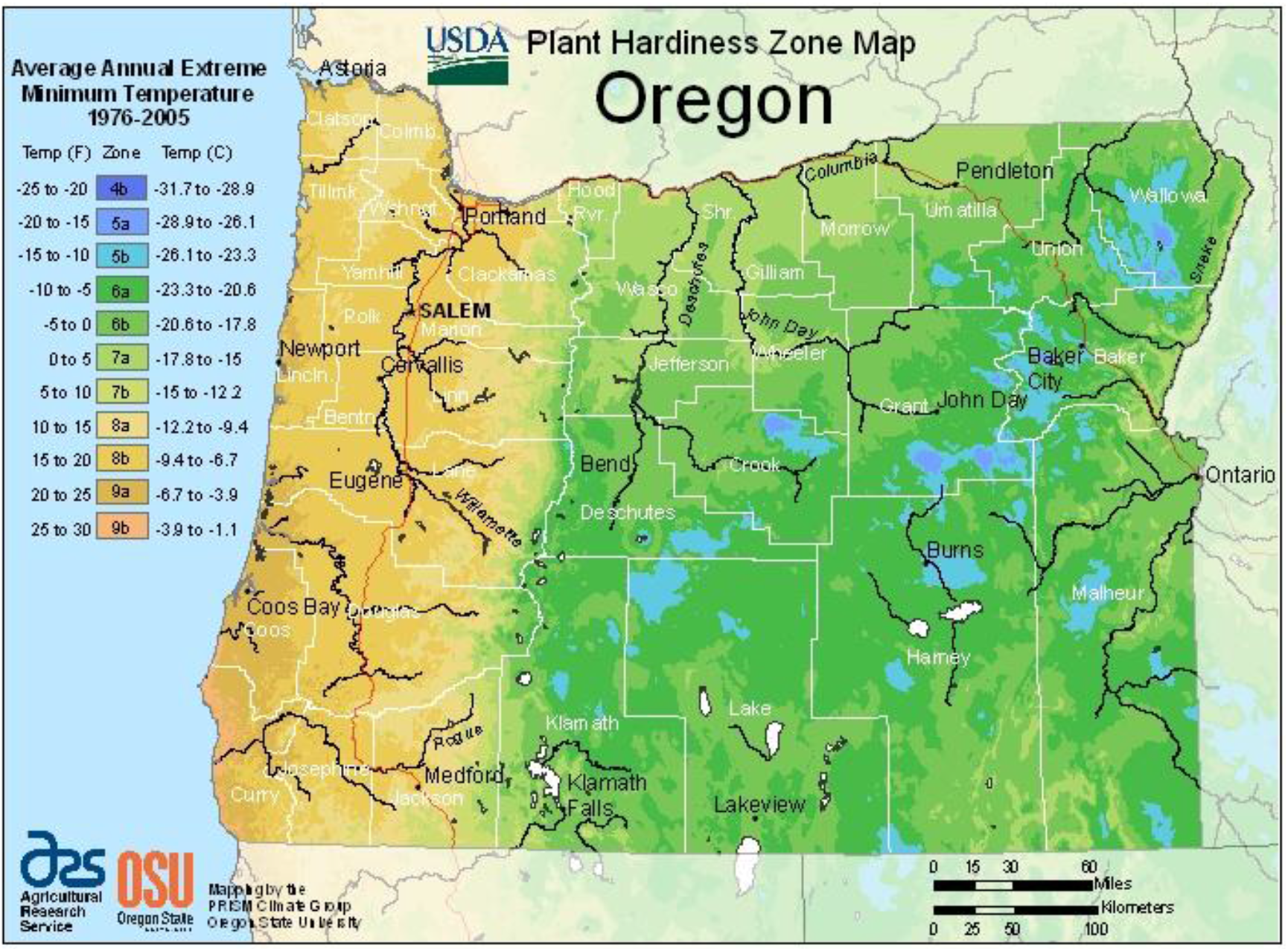
Plant Hardiness Zone (PHZ) Map of Oregon. PHZs are based on the average minimum winter temperature across a 30 year time frame for a region and are used to help growers determine which plants may grow best depending on the zone they inhabit. This risk assessment used PHZs to determine suitable climate where *Vespa mandarinia* may overwinter. Oregon’s PHZs share some of the same PHZs which are present in *V. mandarinia’*s native ranges. These zones include 6A-9B, which include the majority of Oregon, indicating suitable climate for *V. mandarinia* to overwinter in and predate on honeybee populations. (Source: USDA Agricultural Research Service 2020)

The Pacific Northwest also has very dense forest cover. Blackard et al. (2008) estimated that the Pacific Northwest contained the highest densities of forest biomass in the contiguous U.S., with an estimated 9 million ha of forested landcover in Washington, and 13 million ha of forested landcover for Oregon. Considering that *V. mandarinia* prefers to establish and colonize nests within green, herbaceous environments, the Pacific Northwest serves as a suitable region within the U.S. for it to establish and proliferate (Figure 4).

**Figure 4:**
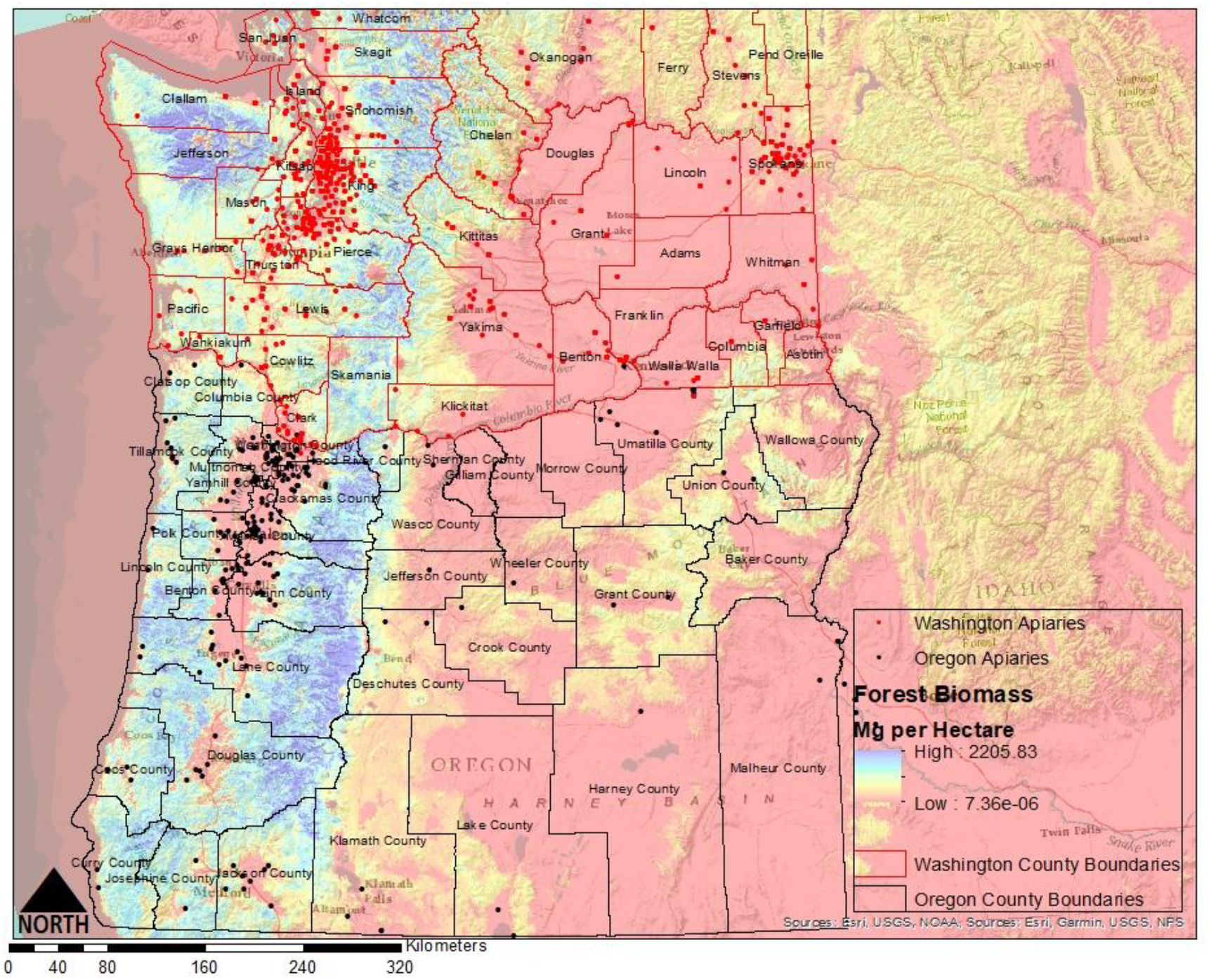
Forest biomass in the Pacific Northwest (Mg/hectare) with apiary distribution in Washington and Oregon. *Vespa mandarinia* prefers to colonize nests in ‘greenspaces’. Considering the density of forest biomass, particularly in the western portions of Washington and Oregon, these regions may serve as suitable habitat for nest establishment. Note the apiary distribution in relation to areas of high forest biomass.

We obtained data on honey bee colony densities and distribution by county for Washington and Oregon based on registered apiaries and number of individual hives of each apiary. The information was then summed for each county for a total number of individual hives in Washington and Oregon, with percentage for each county relative to the total across both states.

For Washington, the results showed that Grant County, Yakima County, and Skagit County comprised the majority of honey bee colonies and apiaries at 40.2%, 12.9%, and 11.0%, respectively, accounting for 64% of the state’s apicultural honey bee populations. The remaining 36% of apiary honey bee populations among Washington’s counties ranged from 0.01% to 3.5% of the state’s total apiary honey bee populations (Table 1).

**Table 1:** Total honey bee hives by county in Washington State and Oregon. Note the high proportions of honey bee hives in Grant, Skagit, and Yakima counties relative to the rest of the state. For Oregon, note Malheur, Linn, Clackamas, Marion, and Yamhill counties relative to the rest of the state. Additionally, these counties also fall within the Plant Hardiness Zones identified to provide suitable climate where *Vespa mandarinia* may overwinter. See Figures 2 and 3 (FOIA Request from Washington Department of Agriculture and Oregon Department of Agriculture).

| **Washington Counties** | **Total hives** | **Oregon Counties** | **Total hives** |
| --- | --- | --- | --- |
| Adams | 100 | Baker | 0 |
| Asotin | 263 | Benton | 4,599 |
| Benton | 42 | Clackamas | 8,026 |
| Chelan | 527 | Clatsop | 30 |
| Clallam | 167 | Columbia | 7 |
| Clark | 2,234 | Coos | 143 |
| Columbia | 1,157 | Crook | 29 |
| Cowlitz | 201 | Curry | 18 |
| Douglas | 3,015 | Deschutes | 23 |
| Ferry | 27 | Douglas | 432 |
| Franklin | 3,009 | Gilliam | 0 |
| Garfield | 508 | Grant | 686 |
| Grant | 36,520 | Harney | 20 |
| Grays Harbor | 94 | Hood River | 2,600 |
| Island | 467 | Jackson | 350 |
| Jefferson | 194 | Jefferson | 12 |
| King | 2,935 | Josephine | 26 |
| Kitsap | 485 | Klamath | 40 |
| Kittitas | 203 | Lake | 15 |
| Klickitat | 29 | Lane | 742 |
| Lewis | 605 | Lincoln | 59 |
| Lincoln | 111 | Linn | 12,457 |
| Mason | 64 | Malheur | 19,100 |
| Okanogan | 544 | Marion | 7,832 |
| Pacific | 37 | Morrow | 0 |
| Pend Oreille | 17 | Multnomah | 1,230 |
| Pierce | 785 | Polk | 2,543 |
| San Juan | 95 | Sherman | 0 |
| Skagit | 10,020 | Tillamook | 112 |
| Skamania | 101 | Umatilla | 3,195 |
| Snohomish | 2,898 | Union | 25 |
| Spokane | 3,623 | Wallowa | 0 |
| Stevens | 3,169 | Wasco | 140 |
| Thurston | 1,305 | Washington | 1,174 |
| Wahkiakum | 36 | Wheeler | 1,000 |
| Walla Walla | 1,915 | Yamhill | 9,767 |
| Whatcom | 1,533 |  |  |
| Whitman | 25 |  |  |
| Yakima | 11,675 |  |  |

For Oregon, the results indicated that Malheur County, Linn County, Yamhill County, Clackamas County, and Marion County accounted for approximately 75% of the state’s honey bee colonies and apicultural honey bee populations, while the residual 29 counties made up the remaining 25% (Table 1).

Based on this information and the Plant Hardiness Zone Maps of Washington and Oregon, and the previously stated habitat suitability for overwintering, the majority of Washington’s and Oregon’s counties fall within these suitable temperature ranges, with the exception of northern Okanogan, Ferry, Stevens, Pend Oreille, and portions of Wallowa, Baker, Grant, Harney, Lake, Malheur, Crook, Deschutes, and Klamath counties (Figure 2 & 3). Although the Plant Hardiness Zones are likely an overgeneralization of suitable habitat for *V. mandarinia* it is nonetheless concerning that most of Washington’s apicultural industry lies within these zones of potentially suitable climate and proximity to areas of suitable habitat for nest colonization (Figures 2, 3, & 4).

## Results and discussion

### Risk characterization

To estimate the risk *V. mandarinia* poses to Washington and Oregon, we used a risk rating and scoring system based on an approach used by Schleier et al. (2007) to rank the relative risk of the following categories and criteria: (1) climate suitability for *V. mandarina* to overwinter based on plant hardiness zones (ideal PHZ score in Tables 1 & 2), (2) habitat suitability to colonize nests in ‘green’ environments which was based on dense forest biomass in the Pacific Northwest, (3) density of apiaries by county, and (4) the proximity of introduction pathways (major port or freight hubs) that may increase the risk of establishment.

**Table 2:** Risk rating table for *Vespa mandarinia* establishment in Washington State. An overall risk rating score (ORS) of 1-4 equals low risk. An ORS of 5-8 equals medium risk, and an ORS of 9-12 equals high risk. Low ORS is highlighted in green, medium in yellow, and high in red.

| County | Ideal Plant<br>Hardiness<br>Zone score | Apiary<br>density score | Dense forest<br>biomass<br>score | Proximity to<br>introduction<br>pathway<br>score | Overall risk<br>rating score<br>(ORS) |
| --- | --- | --- | --- | --- | --- |
| Adams | 2 | 1 | 1 | 2 | 6 |
| Asotin | 2 | 1 | 1 | 2 | 6 |
| Benton | 2 | 1 | 1 | 3 | 7 |
| Chelan | 2 | 1 | 3 | 1 | 7 |
| Clallam | 3 | 1 | 3 | 2 | 9 |
| Clark | 3 | 2 | 3 | 2 | 10 |
| Columbia | 2 | 2 | 2 | 2 | 8 |
| Cowlitz | 3 | 1 | 2 | 3 | 9 |
| Douglas | 2 | 2 | 1 | 1 | 6 |
| Ferry | 1 | 1 | 2 | 1 | 5 |
| Franklin | 2 | 2 | 1 | 3 | 8 |
| Garfield | 2 | 1 | 1 | 1 | 5 |
| Grant | 2 | 3 | 1 | 2 | 8 |
| Grays Harbor | 3 | 1 | 3 | 2 | 9 |
| Island | 3 | 1 | 2 | 1 | 7 |
| Jefferson | 3 | 1 | 3 | 1 | 8 |
| King | 3 | 2 | 2 | 3 | 10 |
| Kitsap | 3 | 1 | 2 | 1 | 7 |
| Kittitas | 2 | 1 | 2 | 1 | 6 |
| Klickitat | 2 | 1 | 1 | 1 | 5 |
| Lewis | 3 | 1 | 3 | 1 | 8 |
| Lincoln | 2 | 1 | 1 | 1 | 5 |
| Mason | 3 | 1 | 3 | 1 | 8 |
| Okanogan | 1 | 1 | 1 | 1 | 4 |
| Pacific | 3 | 1 | 3 | 1 | 8 |
| Pend Oreille | 1 | 1 | 2 | 1 | 5 |
| Pierce | 3 | 1 | 3 | 3 | 10 |
| San Juan | 3 | 1 | 3 | 1 | 8 |
| Skagit | 2 | 3 | 3 | 2 | 10 |
| Skamania | 2 | 1 | 3 | 1 | 7 |
| Snohomish | 3 | 2 | 3 | 3 | 11 |
| Spokane | 2 | 2 | 1 | 3 | 8 |
| Stevens | 2 | 2 | 2 | 1 | 7 |
| Thurston | 3 | 2 | 2 | 1 | 8 |
| Wahkiakum | 3 | 1 | 3 | 1 | 8 |
| Walla Walla | 2 | 2 | 1 | 2 | 7 |
| Whatcom | 2 | 2 | 3 | 2 | 9 |
| Whitman | 2 | 1 | 1 | 1 | 5 |
| Yakima | 2 | 3 | 2 | 1 | 8 |

Those counties in Washington and Oregon with high Plant Hardiness Zone designations received a score of ‘3’. Counties with medium to low PHZ designations received a risk rating of ‘2’ or ‘1’, respectively. Similarly, counties with high densities of forest biomass received a risk rating score of ‘3’, and counties with medium to low forest biomass received risk rating scores of ‘2’ and ‘1’. Counties containing high totals of honey bee hives received a risk rating of ‘3’, while those containing low numbers of honey bee hives relative to the rest of the state received risk ratings of ‘2’ and ‘1’. Lastly, the introduction pathway score was based on major port and freight hubs contained in counties of the Pacific Northwest. If a county contained more than one major port or freight hub, that county received a risk rating of ‘3’. If a county contained 1 major port or freight hub, it received a risk rating score of ‘2’. If a county did not contain a major port or freight hub, it received a risk rating of ‘1’.

The scores were then summed across each risk factor for each county for a total possible overall risk score (ORS) of 12. Those counties which received an overall risk score of 1-4 received a ‘low’ risk rating, while counties that received an overall risk score of 5-8 or 9-12 received a risk rating of ‘medium’ or ‘high’, respectively (Tables 2 & 3). These results are also shown visually (Figure 5). Our results identified 22 counties at a high risk of establishment. Washington contained 9 high risk counties, while Oregon contained 13. For Washington, these counties included Clallam, Clark, Cowlitz, Grays Harbor, King, Pierce, Skagit, Snohomish, and Whatcom. The high risk Oregon counties included Benton, Clackamas, Clatsop, Coos, Douglas, Hood River, Lane, Lincoln, Linn, Marion, Multnomah, Polk, and Yamhill. Of the remaining 53 counties between Washington and Oregon, 48 were found to be at a medium risk of establishment, while only 5 were found to be at a low risk of establishment by *V. mandarinia*.

**Table 3:**
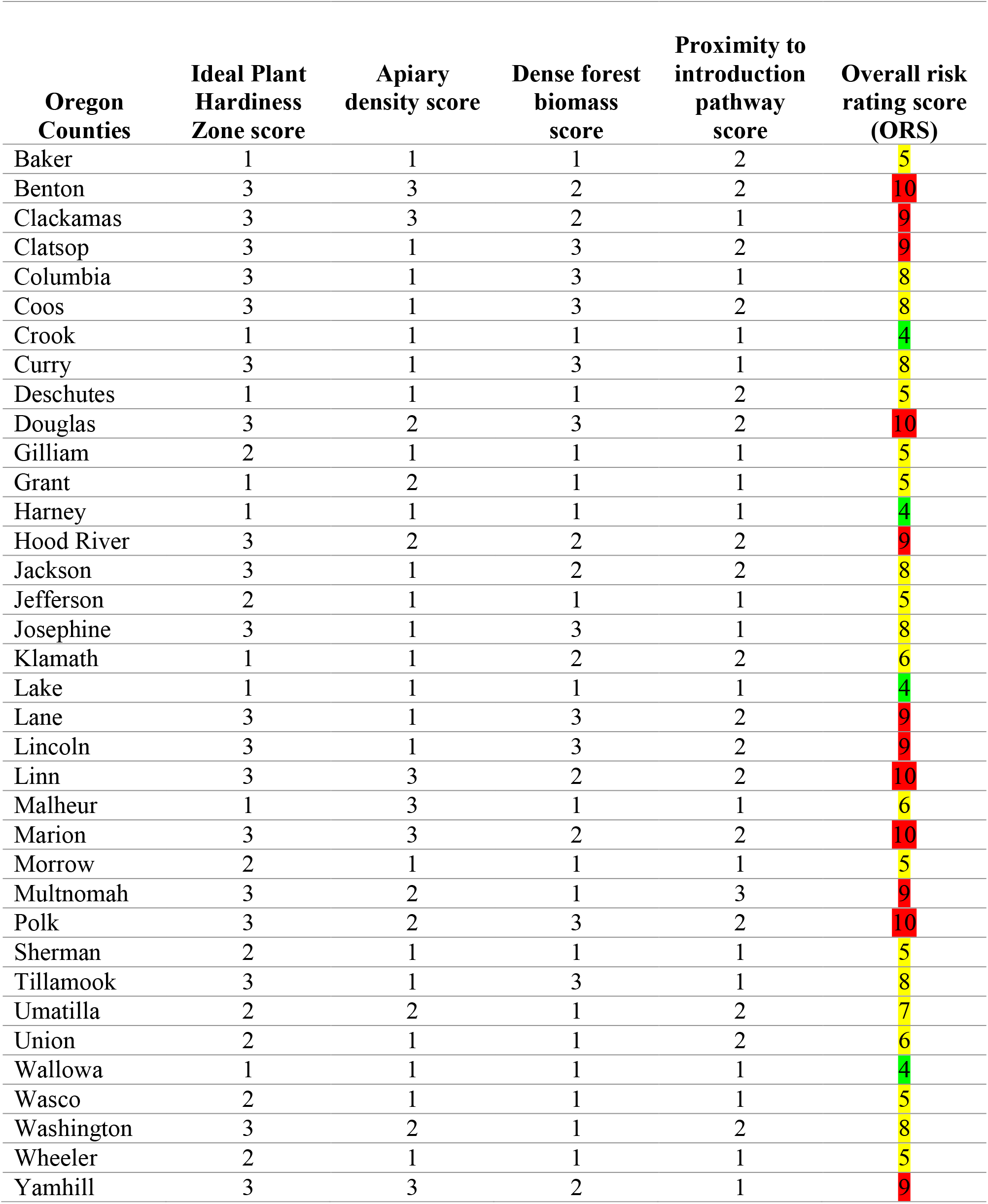
Risk rating table for *Vespa mandarinia* establishment in Oregon. An overall risk rating score (ORS) of 1-4 equals low risk. An ORS of 5-8 equals medium risk, and an ORS of 9-12 equals high risk. Low ORS is highlighted in green, medium in yellow, and high in red.

| Oregon Counties | Ideal Plant Hardiness Zone score | Apiary density score | Dense forest biomass score | Proximity to introduction pathway score | Overall risk rating score (ORS) |
| --- | --- | --- | --- | --- | --- |
| Baker | 1 | 1 | 1 | 2 | 5 |
| Benton | 3 | 3 | 2 | 2 | 10 |
| Clackamas | 3 | 3 | 2 | 1 | 9 |
| Clatsop | 3 | 1 | 3 | 2 | 9 |
| Columbia | 3 | 1 | 3 | 1 | 8 |
| Coos | 3 | 1 | 3 | 2 | 8 |
| Crook | 1 | 1 | 1 | 1 | 4 |
| Curry | 3 | 1 | 3 | 1 | 8 |
| Deschutes | 1 | 1 | 1 | 2 | 5 |
| Douglas | 3 | 2 | 3 | 2 | 10 |
| Gilliam | 2 | 1 | 1 | 1 | 5 |
| Grant | 1 | 2 | 1 | 1 | 5 |
| Harney | 1 | 1 | 1 | 1 | 4 |
| Hood River | 3 | 2 | 2 | 2 | 9 |
| Jackson | 3 | 1 | 2 | 2 | 8 |
| Jefferson | 2 | 1 | 1 | 1 | 5 |
| Josephine | 3 | 1 | 3 | 1 | 8 |
| Klamath | 1 | 1 | 2 | 2 | 6 |
| Lake | 1 | 1 | 1 | 1 | 4 |
| Lane | 3 | 1 | 3 | 2 | 9 |
| Lincoln | 3 | 1 | 3 | 2 | 9 |
| Linn | 3 | 3 | 2 | 2 | 10 |
| Malheur | 1 | 3 | 1 | 1 | 6 |
| Marion | 3 | 3 | 2 | 2 | 10 |
| Morrow | 2 | 1 | 1 | 1 | 5 |
| Multnomah | 3 | 2 | 1 | 3 | 9 |
| Polk | 3 | 2 | 3 | 2 | 10 |
| Sherman | 2 | 1 | 1 | 1 | 5 |
| Tillamook | 3 | 1 | 3 | 1 | 8 |
| Umatilla | 2 | 2 | 1 | 2 | 7 |
| Union | 2 | 1 | 1 | 2 | 6 |
| Wallowa | 1 | 1 | 1 | 1 | 4 |
| Wasco | 2 | 1 | 1 | 1 | 5 |
| Washington | 3 | 2 | 1 | 2 | 8 |
| Wheeler | 2 | 1 | 1 | 1 | 5 |
| Yamhill | 3 | 3 | 2 | 1 | 9 |

**Figure 5:**
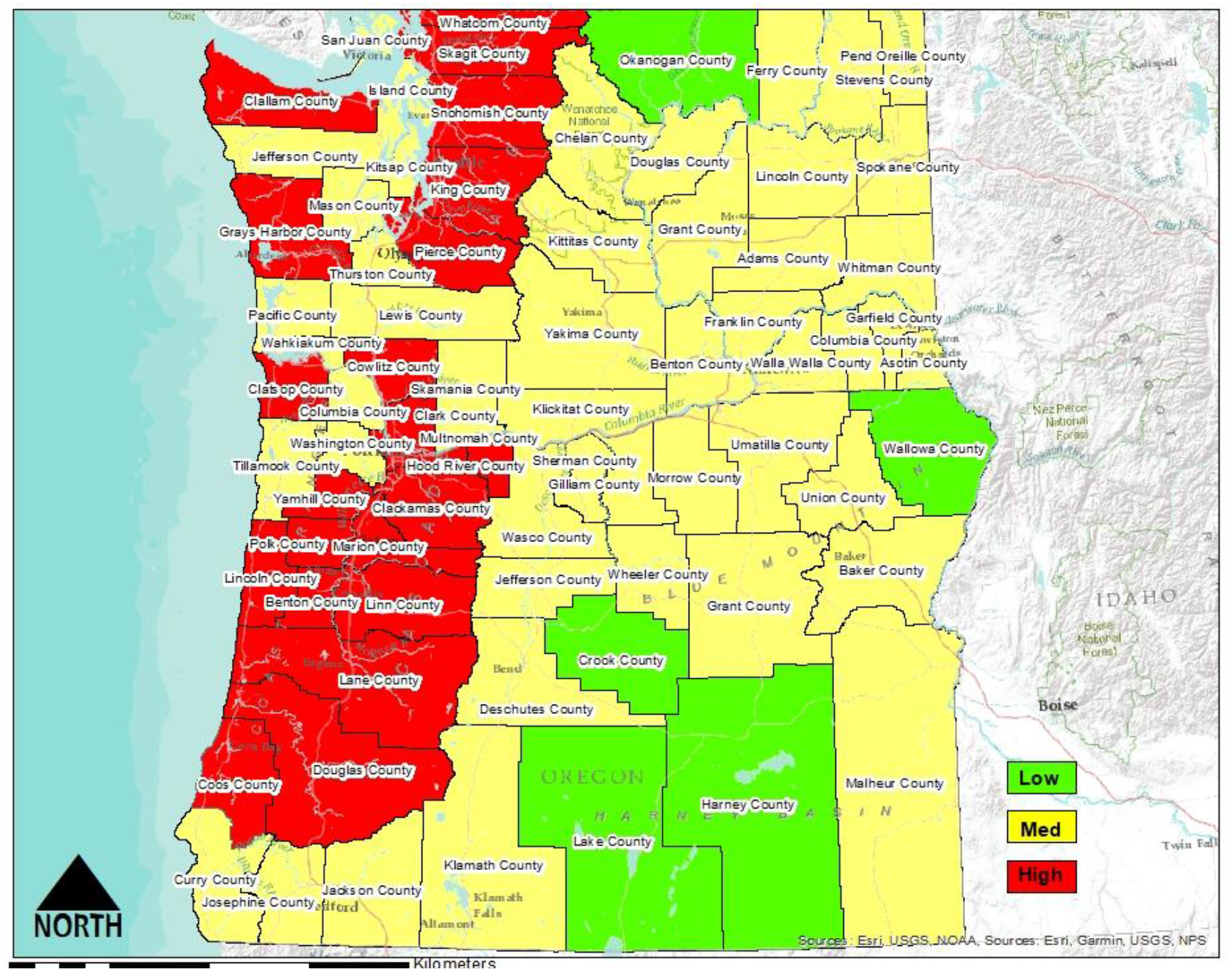
Spatial representation for the overall risk score to each county in the Pacific Northwest for *Vespa mandarinia* establishment.

The results of the risk assessment for the establishment of *V. mandarinia* in the Pacific Northwest suggest that the species may come with serious economic consequences for the region, especially for apiculture and crop-pollinated agriculture sectors. Federal and state regulatory agencies, as well as apicultural and agricultural industries should therefore take immediate action to develop plans and methodologies to prevent further naturalization of *V. mandarinia* in Washington and Oregon, with the ultimate goal of complete eradication of the species.

### Uncertainty analysis

Our risk assessment had the goal of delineating the biological requirements of *V. mandarinia* and analyzing its known ecological relationships with the environments of the Pacific Northwest to estimate the establishment risk of the hornet on a county-by-county basis. The results suggest a number of potential high, medium, and low risk factors that may aid in its establishment in the Pacific Northwest.

However, this risk assessment used a tier-1 or screening level approach, which is usually employed when there is a lack of quantitative or spatial data. Considering that *V. mandarinia* was just recently introduced into the Pacific Northwest, there are few data concerning the species’ current distribution within Washington or elsewhere in the Pacific Northwest. Moreover, there is very little published literature on the species, with most of the published research dating back to the 1970s through the 1990s. Furthermore, although there has been somewhat of a recent resurgence in the literature on the species due to the recent introduction of the species into North America, most of this research also relied heavily on the same aforementioned research that was published decades ago.

With this scarcity of data and lack of knowledge concerning the species’ ecological relationships and distribution in the Pacific Northwest, our risk assessment relied on only a few of the biotic and abiotic requirements which may sustain or hamper establishment success of the hornet. For example, the use of Plant Hardiness Zones to delineate habitat suitability for *V. mandarinia* in the Pacific Northwest likely considerably overestimates the areas in which the species could survive its overwintering period and establish new colonies the following year. Instead of using Plant Hardiness Zones, other approaches may be more accurate and therefore informative. For example, CLIMEX or Maxent species distribution modeling may prove useful as more worldwide occurrence data and documented physiological tolerances become available (Kumar et al. 2016, Kriticos et al. 2015). Accordingly, more detailed research regarding the ecological relationships and life cycle of *V. mandarinia* in the Pacific Northwest needs to be undertaken to form a more complete picture of what can actually be defined as suitable habitat within this region.

Similarly, the honey bee hive distribution and density data were based solely on hives which were managed by registered apiaries and did not take into account hives which may be managed by unregistered beekeepers. In addition, although the risk to apiaries may be easier to estimate, our risk assessment was not able to assess the risk to the pollination services provided by wild bee populations considering that to our knowledge no data exist on estimates of wild bee populations in the Pacific Northwest.

In addition, although the use of risk-rating systems in qualitative or semi-qualitative risk assessments aid in producing simple categorizations of risk based on supporting reasoning and documentation (Cox et al. 2005), they are not without limitation. Risk rating systems often lack the confidence to accurately discern between quantitatively small and quantitatively large risks. This can result in errors such as the assignment of higher risk ratings to either a particular, or multiple, risk situations which may in reality actually differ quantitatively by orders of magnitude (Cox et al. 2005).

Furthermore, our risk characterization’s reliance on estimating risk on a county by county basis is a coarse scale, which likely results in an over or under estimation of the actual risk in a given area. Future assessments should focus on estimating risk in the region at a finer scale.

## Acknowledgements

We thank T. Sterling (Montana State University) for reviewing an earlier version of the manuscript. This research was funded in part by the Montana Agriculture Experiment Station, and Montana State University. This material is based on work that is supported by the National Institute of Food and Agriculture, Hatch Multistate Project No. W-4045.

## Abbreviations

*A. cerana japonica*: *Apis cerana japonica*
*A. mellifera*: *Apis mellifera*
ORS: Overall Risk Score
PHZ: Plant Hardiness Zone
*V. mandarinia*: *Vespa mandarinia*

## Disclaimer

The findings and conclusions in this publication are those of the authors and should not be construed to represent any official USDA or U.S. Government determination or policy.

